# Characterization of Dr20/22: Molecular Insights and Antibody Response in Naturally Infected Dogs with *Dirofilaria repens*

**DOI:** 10.1101/2023.09.15.557890

**Authors:** Mateusz Pękacz, Katarzyna Basałaj, Daniel Młocicki, Maciej Kamaszewski, Elena Carretón, Rodrigo Morchón, Marcin Wiśniewski, Anna Zawistowska-Deniziak

**Affiliations:** Division of Parasitology, Department of Preclinical Sciences, Faculty of Veterinary Medicine, Warsaw University of Life Sciences-SGGW, 02-786 Warsaw, Poland; Museum and Institute of Zoology, Polish Academy of Sciences, 00-679 Warsaw, Poland; Department of General Biology and Parasitology, Medical University of Warsaw, 02-004 Warsaw, Poland; Department of Ichthyology and Biotechnology in Aquaculture, Institute of Animal Sciences, Warsaw University of Life Sciences - SGGW, 02-786 Warsaw, Poland; Internal Medicine, Faculty of Veterinary Medicine, University of Las Palmas de Gran Canaria, Campus Arucas, Arucas, 35413 Las Palmas, Spain; Zoonotic Diseases and One Health Group, Faculty of Pharmacy, University of Salamanca, Campus Miguel Unamuno, 37007 Salamanca, Spain; Department of Immunology, Institute of Functional Biology and Ecology, Faculty of Biology, University of Warsaw, 02-095 Warsaw, Poland

**Author notes:** These authors contributed equally to this work (Marcin Wiśniewski, Anna Zawistowska-Deniziak).

**Keywords:** dirofilariasis, diagnostics, *alt*, spliced leader trans-splicing

## Abstract

Subcutaneous dirofilariasis, caused by the parasitic nematode Dirofilaria repens, is a growing concern in Europe, affecting both dogs and humans. This study focused on D. repens Dr20/22, a protein encoded by an alt (abundant larval transcript) gene family. While well-documented in L3 larvae of other filariae species, this gene family had not been explored in dirofilariasis. The research involved cloning Dr20/22 cDNA, molecular characterization, and evaluating its potential application in the diagnosis of dirofilariasis. Although Real-Time analysis revealed mRNA expression in both adult worms and microfilariae, the native protein remained undetected in lysates from both developmental stages. This suggests the protein’s specificity for L3 larvae and may be related to a process called SLTS (spliced leader trans-splicing), contributing to stage-specific gene expression. The antigen’s apparent specificity for invasive larvae makes it a strong candidate for an early dirofilariasis marker. ELISA tests using sera from infected and healthy dogs detected specific IgG antibody responses in both groups, indicating that specific IgG in healthy dog sera may contribute to dirofilariasis immunity. Although further research is necessary, the molecular and immunological characterization of Dr20/22 can enhance our understanding of host-parasite interactions and provide insights into the mechanisms that facilitate immune system evasion.

## 1. Introduction

Subcutaneous dirofilariasis is a vector-borne disease caused by a parasitic nematode Dirofilaria repens. Owing to climate changes and anthropogenic activities, dirofilariasis has expanded its distribution across nearly all European countries [1]. Consequently, it has emerged as one of the most rapidly spreading parasitoses in the Old World. While the primary hosts of the disease are carnivores, notably dogs, it exhibits considerable zoonotic potential, thereby constituting a significant concern in both medical and veterinary fields. However, due to the predominantly asymptomatic or mildly symptomatic nature of D. repens infections [2], this zoonosis has often been overlooked, particularly when compared to other filariasis such as pulmonary dirofilariasis or lymphatic filariasis. As a result, research efforts aimed at mitigating invasions, such as early infection detection or development of potential vaccines, have been substantially impeded, given the limited understanding of the parasite’s biology and the molecular interactions it established with the hosts.

Throughout the protracted co-evolution between hosts and parasites, helminths have evolved diverse strategies to evade host immune responses at various stages of their life cycle [3–6]. Notably, the filariae L3 larvae adeptly evade and down-modulate the host’s immune system within the cutaneous tissues, playing a pivotal role in both the parasite’s establishment and the development of host immunity [7]. These immunomodulatory properties are primarily attributed to the surface and secreted antigens displayed by the larvae. As the initial molecules that engage the definitive host’s immune system, these antigens hold promise as potential early markers of infection. Moreover, the presence of antibodies against L3 larvae is hypothesized to suppress larval development and correlate with resistance to new infections [8,9]. However, despite the critical roles, the specific molecules underlying immune modulation in D. repens remain elusive. Thus, an in-depth investigation into stage-specific antigens, along with their molecular and immunological characterization, may offer novel insights into the development of prospective vaccines or diagnostic tools.

In light of this, the present study focused on a singular L3-specific protein known as D. repens 20/22 kDa (Dr20/22), which was produced in recombinant form. Dr20/22 is encoded by a gene belonging to the alt (abundant larval transcript) family, a group well documented in numerous filarial nematodes including Brugia malayi [10,11], Onchocerca volvulus [12], Acanthocheilonema vitae [13], Wuchereria bancrofti [14] and Dirofilaria immitis [15,16]. The ALTs being abundantly expressed in invasive larvae and devoid of known homologues in mammals, represent highly attractive vaccine candidates for filarial infections. While the precise biological function of these antigens remains obscure, extensive research has been conducted to explore their prophylactic potential, particularly in human lymphatic filariasis caused by B. malayi and W. bancrofti [11,14,17–24]. Intriguingly, however, no investigations have yet delved into this antigen in the context of subcutaneous dirofilariasis. Thus, the D. repens homologue of ALT might hold central importance in the invasion process, and its comprehensive molecular and immunological characterization could represent a critical milestone in advancing our understanding of the intricate host-parasite interactions.

This study entailed the cloning of cDNA encoding the D. repens homologue of ALT (Dr20/22), followed by its meticulous molecular characterization. Subsequently, the recombinant antigen was evaluated for its potential application as an early diagnostic marker for subcutaneous dirofilariasis and used to discern antibody responses in dogs naturally infected with D. repens in comparison to non-infected individuals.

## 2. Results

### 2.1. Cloning and Characterization of cDNA and gDNA of the Dr20/22 Gene in D. repens

The cDNA cloning of the dr20/22 gene yielded a 417 bp product. The complete coding sequence of the *D. repens alt* gene homologue in its adult stage was submitted to GenBank under accession number MN706526.1. No *D. repens*-specific SL sequence was found in either the adult stage or the microfilariae stage cDNA. Additionally, the gDNA cloning of almost the entire gene (from the start codon to the stop codon) resulted in a 1,048 bp product (Supplementary File S1).

### 2.2. Comparative Analysis of Nucleotide and Amino Acid Sequences of ALT cDNAs From Related Filariae Reveals Phylogenetic Relationships and Structural Characteristics of Dr20/22 Protein

Comparison of nucleotide sequences of ALT cDNAs from related filariae using Blast showed 82.12 % identity to *D. immitis* (U29459.1), 76.19 % to *Loa loa* (XM_003148107), 60.92 % to *O. volvulus* (U29576.1), 78.52 % to *A. vitae* (U47545.2), 77.29 % to *W. bancrofti* (AF285860.1), and 72.98 % to *B. malayi* (U84723.1). A comparison of cDNA and gDNA encoding Dr20/22 revealed that the gene consists of 4 exons (1-78 bp; 296-422 bp; 665-755 bp; 928-1,048 bp) and 3 introns (Supplementary File S1).

Comparison of amino acid sequences showed 75.69 % (MCP9263779.1) and 72.85 % (AAC47031.1) identity to *D. immitis*, 55.48 % to A. vitae (AAB03902.2), 46.81 % to *O. volvulus* (AAA84910.1), 64.10 % to *L. loa* (XP_003148155.1), 56.62 % to *W. bancrofti* (EJW81953.1), and 53.68 % to *B. malayi* (AAB41884.1).

Analysis of the putative amino acid sequence revealed the presence of a 21 amino acid signal peptide, followed by a protein with a theoretical molecular mass of 13.82 kDa and pI 4.93. NetOGlyc and NetNGlyc biotools determined six potential O-glycosylation sites (in positions 27, 29, 33, 36, 53, 60) and no N-glycosylation sites. NetPhos revealed 18 potential sites of phosphorylation: serine (23, 27, 29, 33, 36, 67, 97, 107, 118, 125, 128), threonine (53, 60), tyrosine (37, 41, 58, 81, 126). InterProScan identified a Chromadorea ALT domain (IPR008451) in position 62-135 characteristic of a vast family of filariae. The Phyre2 program constructed the most probable 3D structure based on bee-venom phospholipase A2 from Apis mellifera (PMID: 2274788) but the confidence of the model was estimated at only 3.8 % (Figure 1).

**Figure 1.**
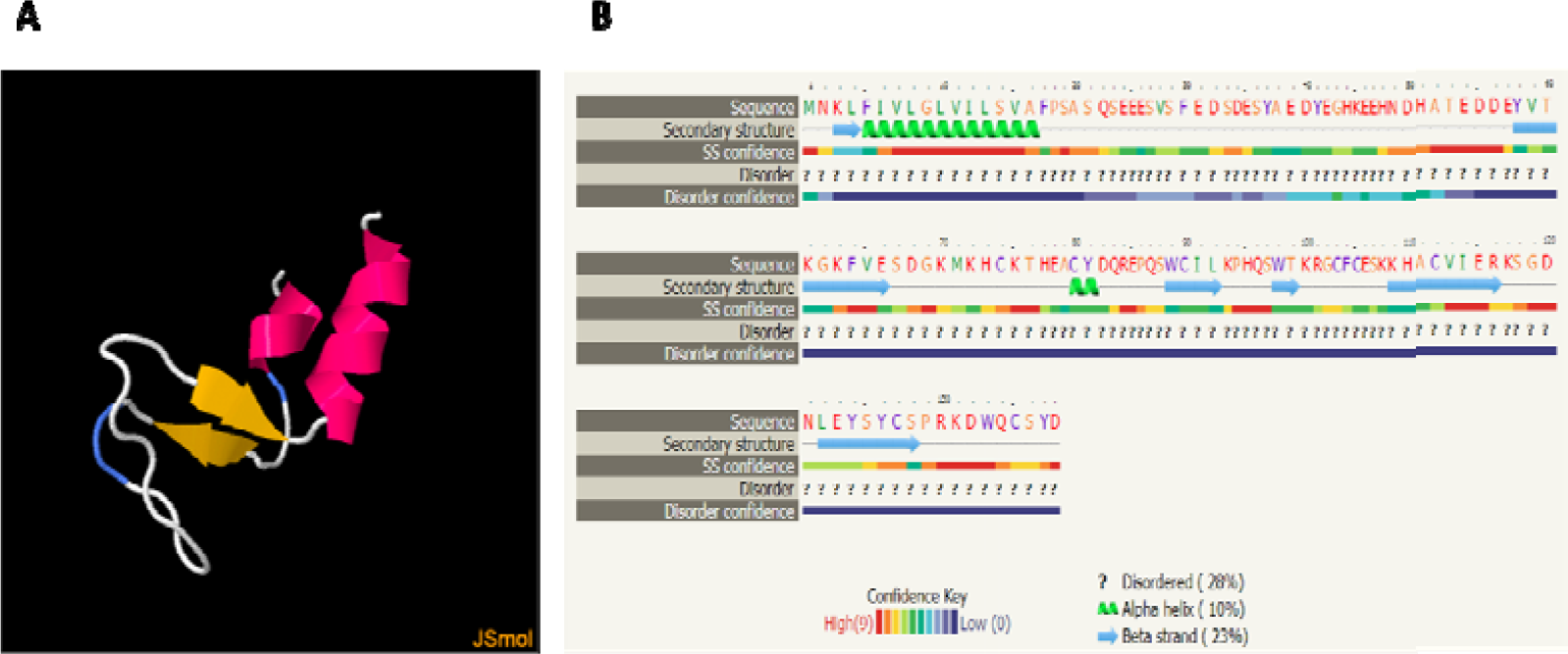
Probable three-dimensional structure of the *D. repens* ALT protein (Dr20/22) constructed in Phyre2 (A) and its secondary structure prediction (B).

### 2.3. Real-Time PCR Analysis of D. repens alt Gene Expression

Gene expression analysis showed 3.32 times higher expression levels of the D. repens alt gene in the adult stage than in microfilariae (Figure 2).

**Figure 2.**
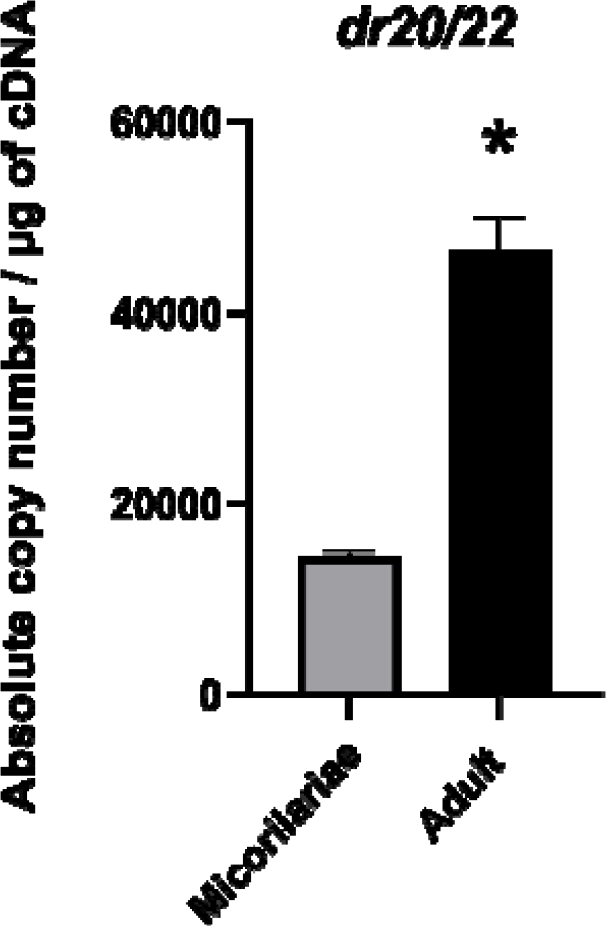
Analysis of the expression levels of *dr20/22* in microfilariae and the adult stage of *D. repens*. The result shows the number of gene copies per µg of cDNA. The bars show mean ± SEM. The statistically significant differences between examined groups are marked with asterisk : \****p*** < 0.05.

### 2.4. Expression and Characterization of Dr20/22 Recombinant Protein

Dr20/22 was efficiently expressed in BMMY medium, yielding a high amount of the recombinant protein. Subsequent SDS-PAGE and Western blot analyses displayed the purified protein as a double band within the 20-22 kDa range (Figure 3). Glycoprotein staining did not indicate the presence of any glycan residues (Figure 4). Interestingly, when the gel was overloaded with the protein, smaller fragments of the protein were detected, implying a propensity of the protein to undergo collapse. This observation was further confirmed in subsequent Western blot analyses using frozen protein.

**Figure 3.**
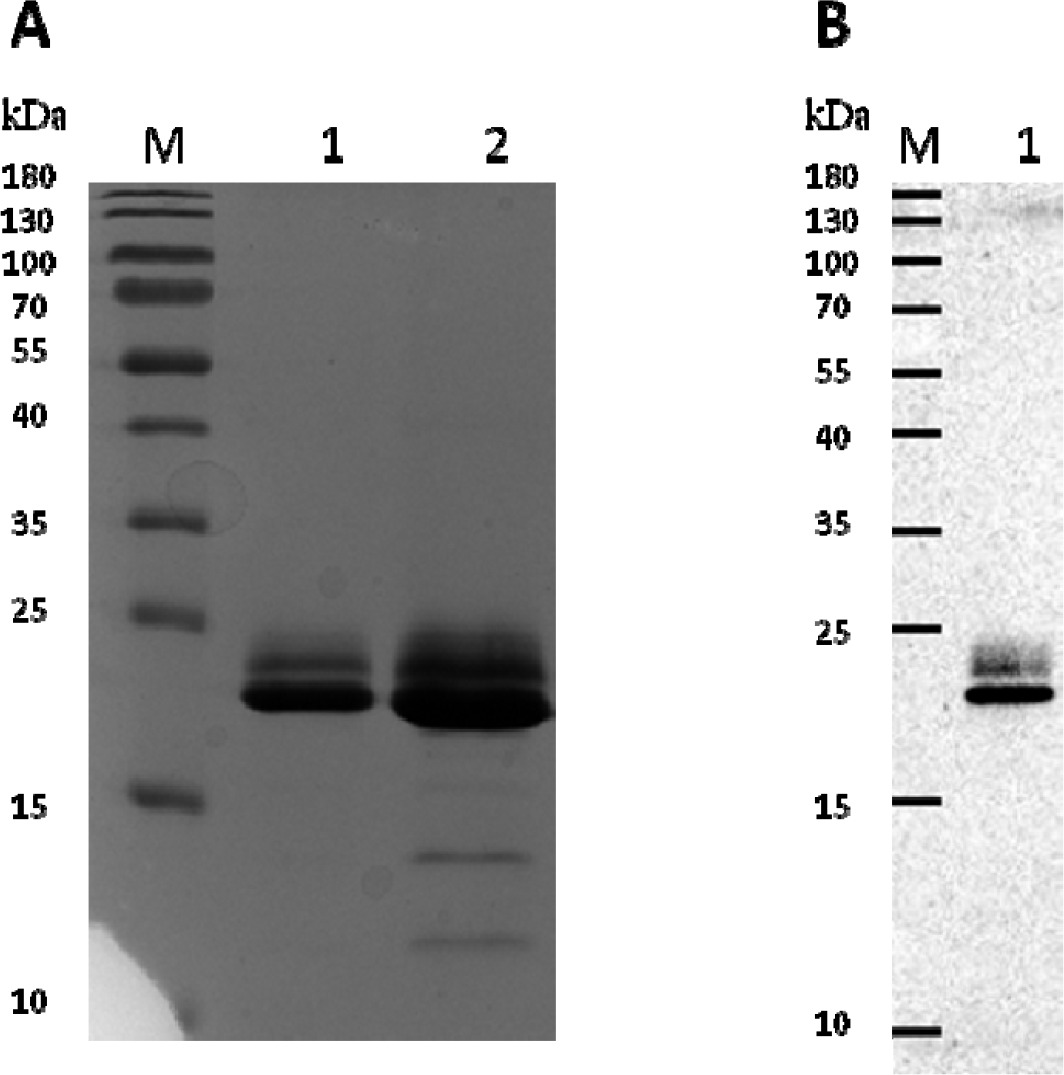
SDS PAGE (A) and Western blot (B) analysis of purified Dr20/22. Lane M - molecular weight marker (PageRuler™ Prestained Protein Ladder, 10 to 180 kDa, Thermo Scientific™); lane 1A - 5 μg of the rDr20/22; lane 2A - 15 μg of the rDr20/22; lane 1B - 0.25 μg of the rDr20/22.

**Figure 4.**
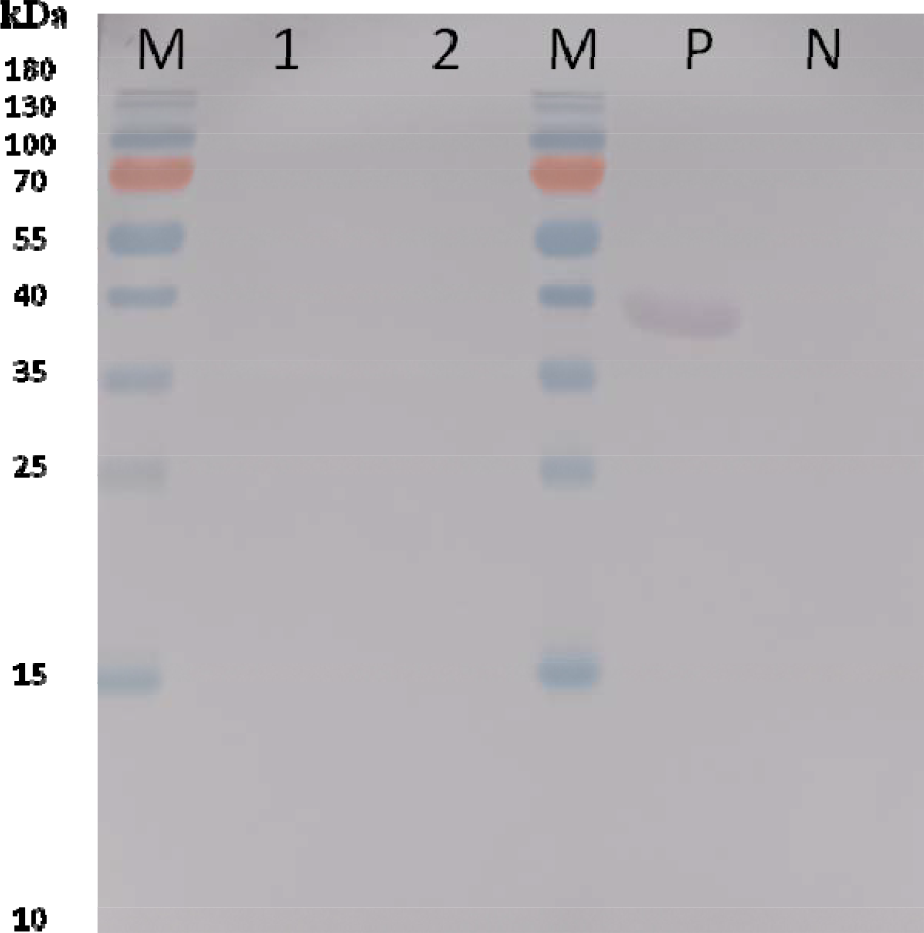
Staining of glycosylated proteins. Lane M - weight marker (PageRuler™ Prestained Pro- tein Ladder, 10 to 180 kDa, Thermo Scientific™); lane 1 - 5 μg of the rDr20/22; lane 2 - 10 μg of the rDr20/22-; P - positive control; N - negative control.

### 2.5. PLA2 Assay Has Shown No Activity of the Dr20/22

As PLA2 activity was not detected, we conducted tests to explore the protein stability and the potential impact of freezing on antigen activity. Both fresh protein (used immediately after purification) and protein subjected to a freeze-thaw cycle were examined. The rationale behind this testing approach was to ensure that if the protein did possess PLA2 activity, it could be assessed for any potential loss or changes in activity resulting from the freezing process.

### 2.6. Lack of Dr20/22 Detection in D. repens Lysates from Adult and Microfilariae Stages

The Western blot analysis confirmed the presence of antibodies targeting the recombinant Dr20/22 epitopes in the anti-Dr20/22 serum, while no specific antibodies were detected in the serum of non-immunized mice. It was previously mentioned that the freeze-thaw cycle resulted in the observation of smaller protein fragments (Figure 5).

**Figure 5.**
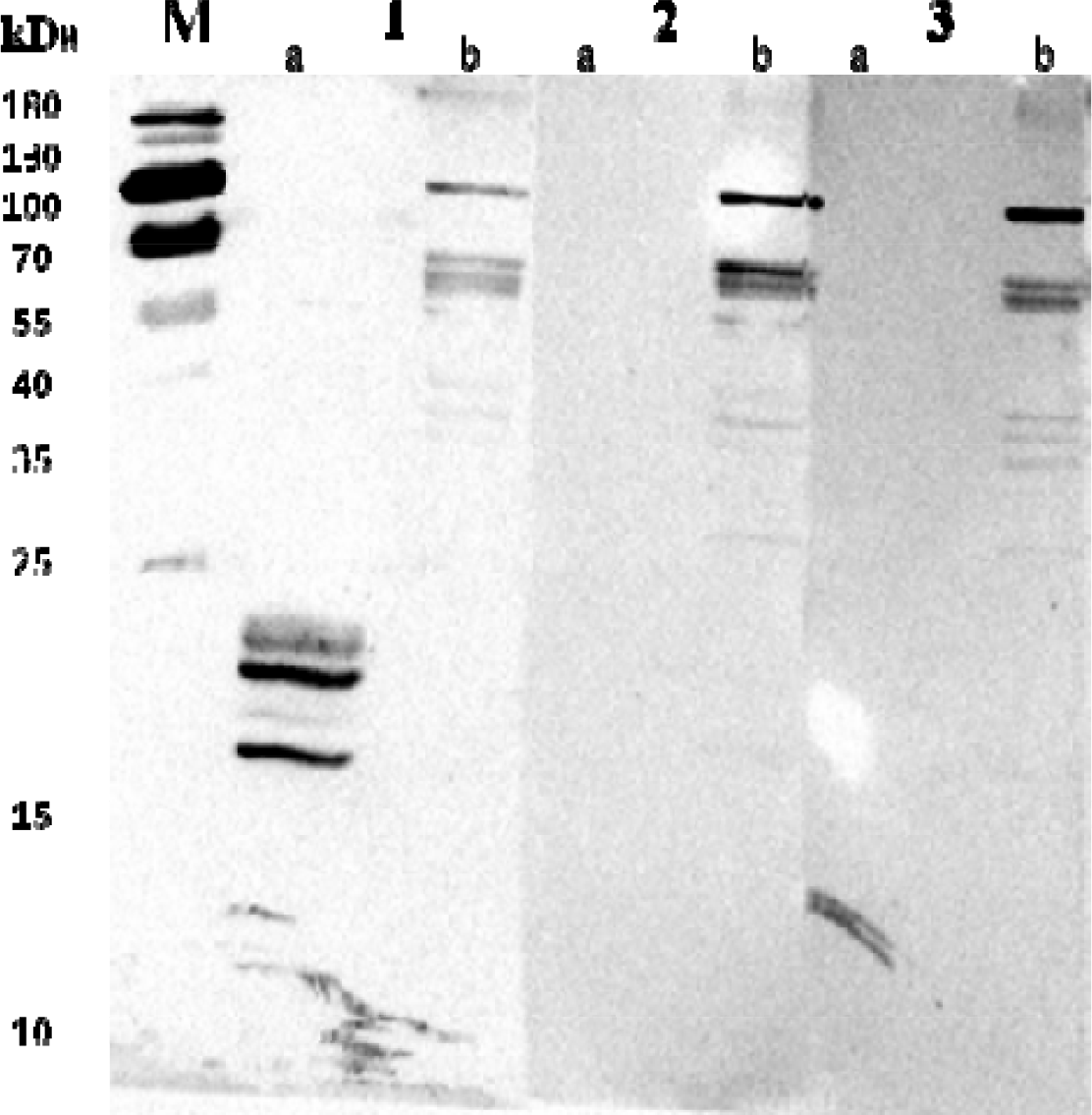
Western blot analysis of the specificity of anti-Dr20/22 antibodies to recombinant Dr20/22 (a) and native protein (b) in adult D. repens homogenate using mouse anti-Dr20/22 serum (1), and sera from non-immunized mice (2 and 3). Lane M - molecular weight marker (PageRuler™ Prestained Protein Ladder, 10 to 180 kDa, Thermo Scientific™).

Native Dr20/22 was not detected in any of the tested D. repens lysates, including adult female worm, E/S fraction, and microfilariae. Interestingly, some cross-reactions were observed between the adult worm extract and sera from both immunized and negative mice (Figure 5). However, specific localization of the protein in cross-sections of the adult female worm was not achieved (Figure 6). The anti-Dr20/22 serum showed a strong signal almost throughout the entire histological section, while a weak signal was also observed with the serum from non-immunized mice, suggesting the possibility of cross-reactions (Figure 7).

**Figure 6.**
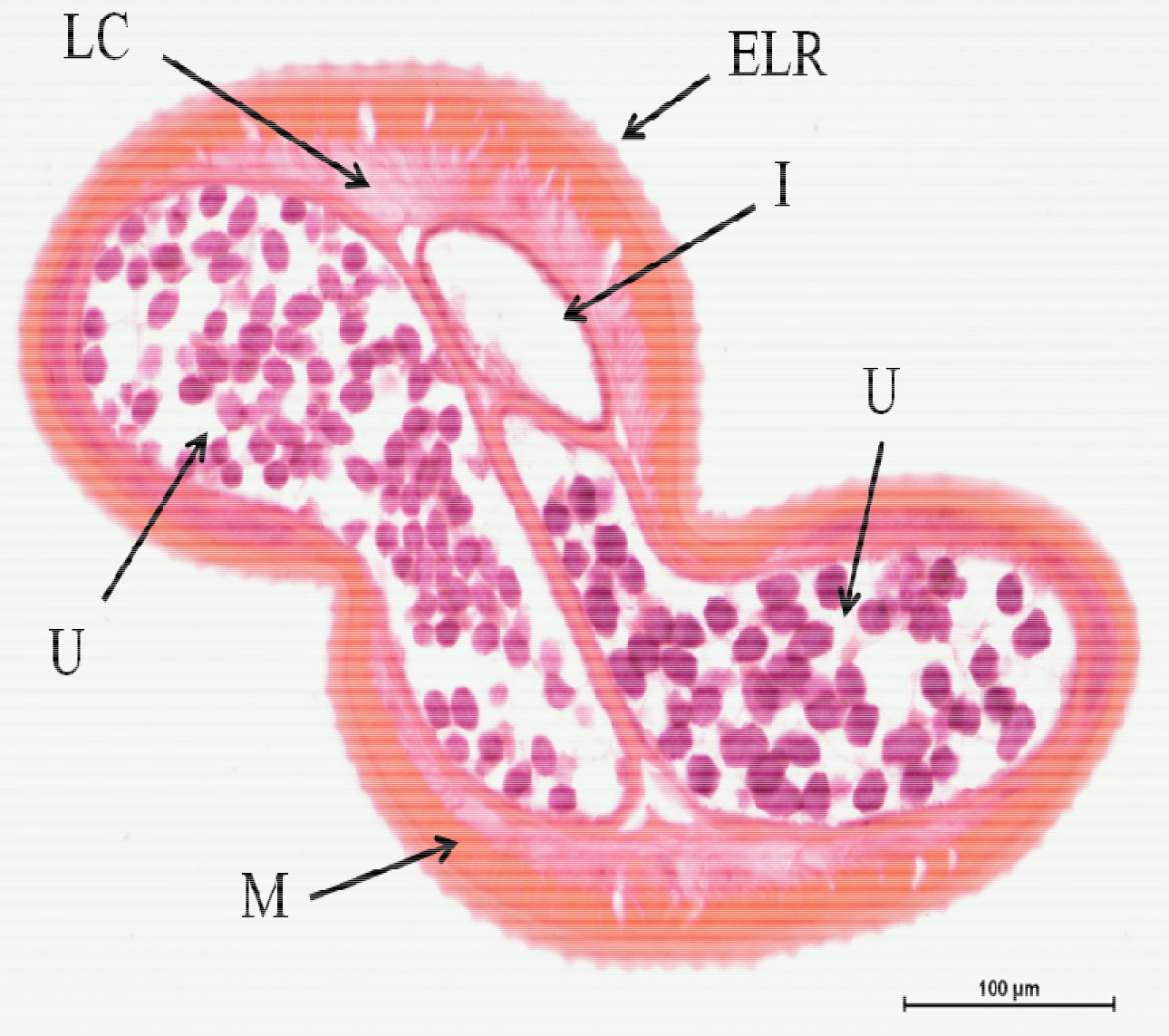
Morphological details of a D. repens female transverse section stained with H&E. ELR - external longitudinal ridges; CL - cuticular layer; LC - lateral chord; M - musculature; I - intestine; U - uterus containing microfilariae.

**Figure 7.**
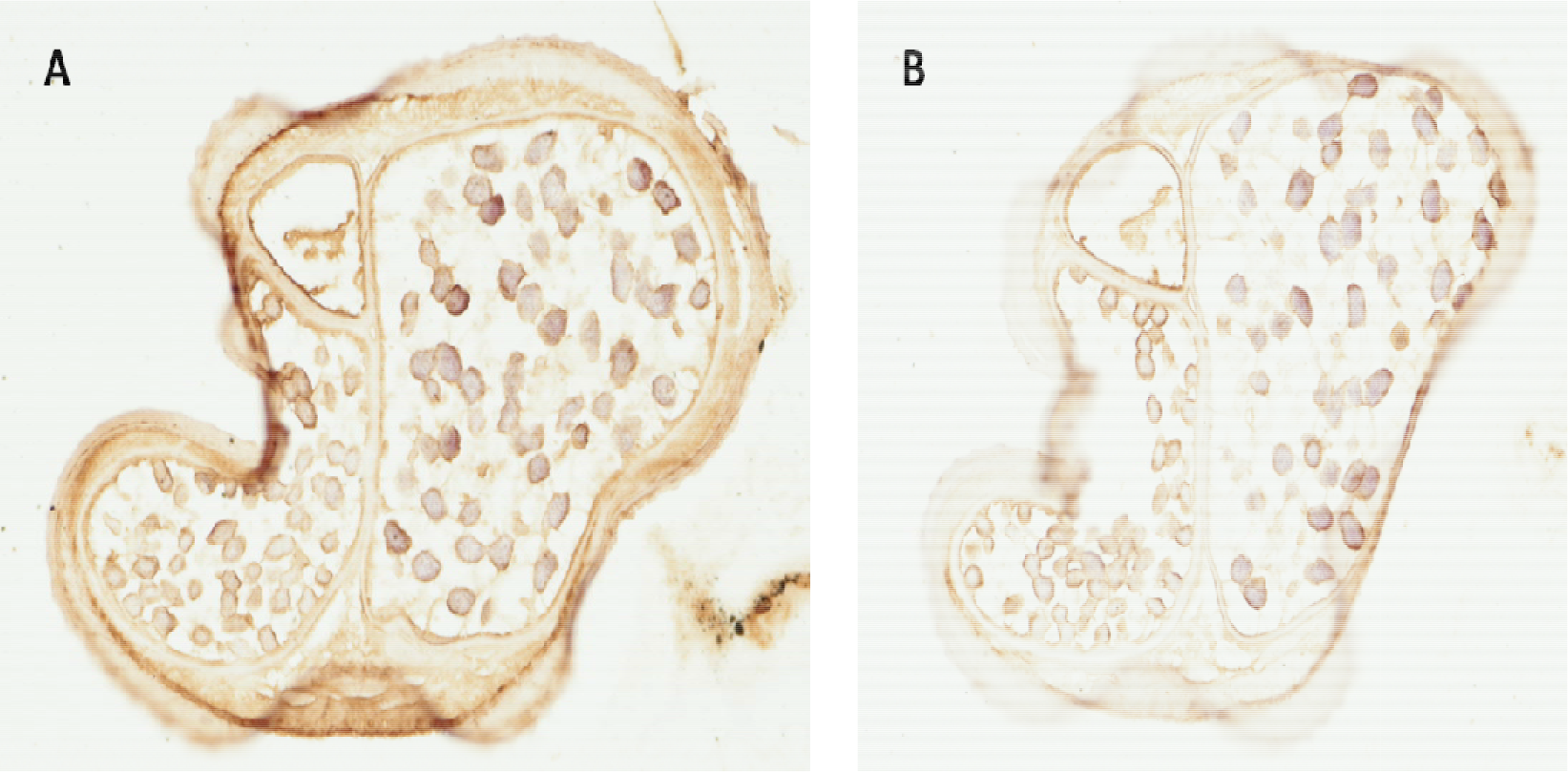
Immunohistochemistry staining of a D. repens female transverse sections with anti-Dr20/22 serum (A) and non-immunized mouse serum (B).

### 2.7. Recognition of Dr20/22 by Antibodies in Sera From Naturally Infected Dogs with D. repens

Our Western blot analysis revealed the presence of specific IgG antibodies against Dr20/22 in sera from dogs infected with *D. repens*, while no such antibodies were detected in dogs infected with *D. immitis* (Figure 8). Interestingly, in the Dot blot analysis, majority of *D. immitis* sera recognized the protein (Figure 9). These results suggest that the antigenicity of Dr20/22 may be influenced by its conformation, with potential variations in response between its native and denatured states.

**Figure 8.**
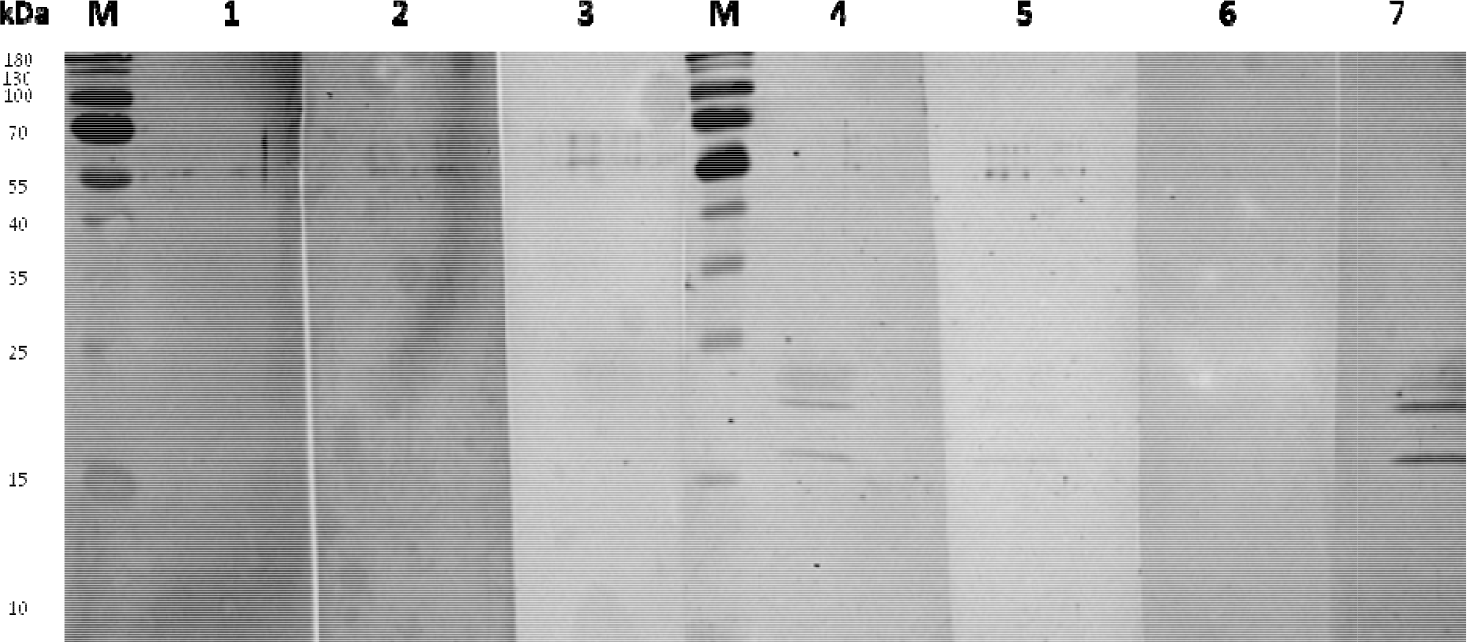
Western blot analysis of the presence of IgG specific to rDr20/22 in sera from dogs infected with D. immitis (1-3), D. repens (4, 5, 7) and negative dog (6). Lane M - molecular weight marker (PageRuler™ Prestained Protein Ladder, 10 to 180 kDa, Thermo Scientific™).

**Figure 9.**
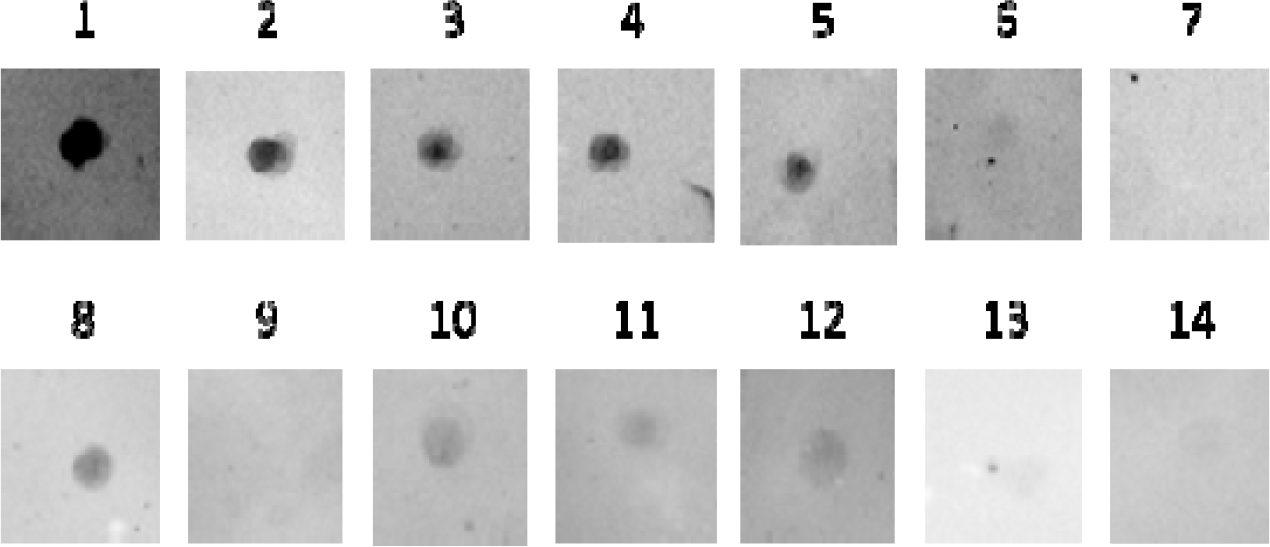
Dot blot analysis of the presence of IgG specific to non-denaturated rDr20/22 in sera from dogs infected with D. repens (1-5), D. immitis (8-12), and negative dogs (6, 7, 13, 14).

### 2.8. ELISA with DrSA Enables a Specific Diagnosis of the Infection but Presents Cross-Rea tions with D. immitis

In this study, we analyzed sera from 850 dogs, comprising 174 with active microfilaremia and 676 potentially negative cases, using DrSA ELISA. Since molecular testing was not feasible for all samples, we compared them against a calculated c value. This value was determined based on 35 dogs known to be negative in Knott/ t-off PCR and ELISA, as described in our previous study [25]. Dogs with IgG levels above the cut-off were classified as amicrofilaremic. Interestingly, we observed “low responders”, a subgroup of dogs with active microfilaremia but IgG levels below the cut-off. To categorize the dogs, we divided them into three groups: microfilaremic (Mf+), amicrofilaremic (Mf-), and negative (Neg) **(Figure 10)**. We then explored the IgM class in the dogs based on this classification, but no significant differences between infected and non-infected groups were found (Figure 11). Considering that some dogs classified as negative in the IgG class might be in the early infection phase and only have increased IgM levels, or they could be occult “low responders”, we decided to compare microfilaremic dogs with “true negative” dogs (with low levels of both IgG and IgM). Interestingly, we observed a higher level of IgM in the infected group **(Figure 12)**. Despite the sensitivity of DrSA ELISA in diagnosing dirofilariasis, we sought to determine if it could differentiate between *D. repens* and *D. immitis* infections, given their genetic closeness. To investigate this, we tested the cross-reactivity between IgG and IgM antibodies from dogs infected with either *D. repens* or *D. immitis* using somatic antigens from adult worms of both species (DrSA vs. DiSA) **(Figure 13)**.

**Figure 10.**
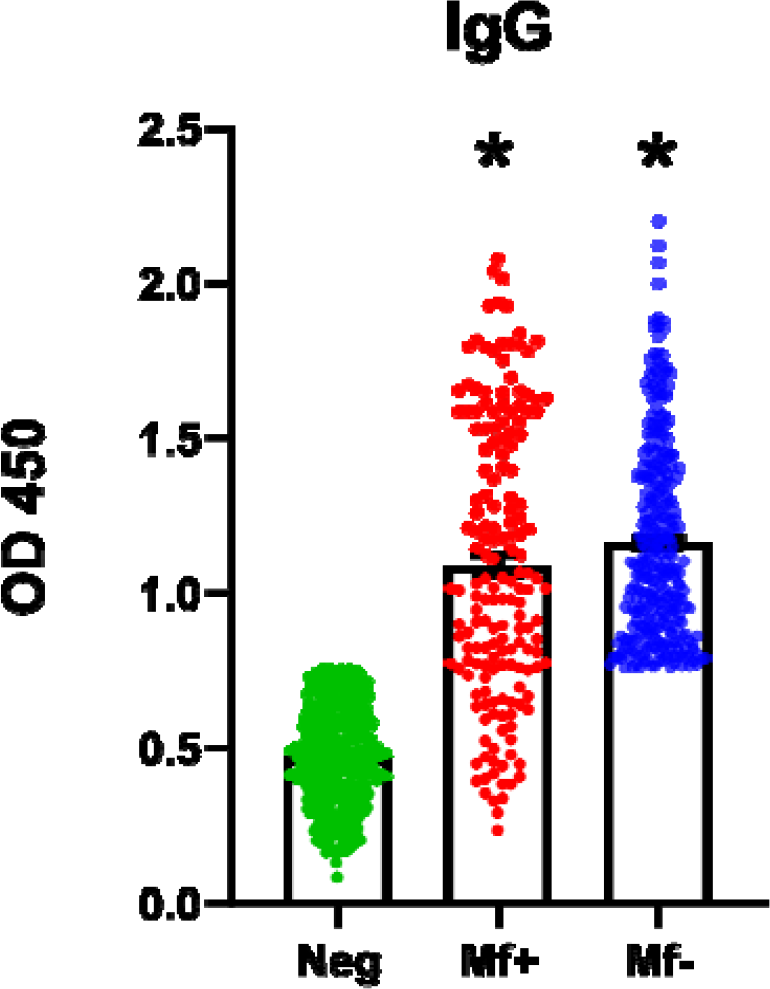
D. repens somatic antigen (DrSA) ELISA enables to detect infections in microfilaremic and amicrofilaremic dogs. Sera from potentially negative dogs (n = 676) were compared against calculated cuf-off value determined based on 35 dogs known to be molecularly negative. Finally, dogs were categorized into three groups: microfilaremic (red; N = 174), amicrofilaremic with IgG level above cut-off value (blue; N = 297) and negative - with IgG level below the cut-off value (green; N = 379). The IgG level was analyzed in dogs sera against DrSA. Plates were coated with 2.5 µg/ml of DrSA in carbonate buffer. The blocking step was performed with 0.1 M NaHCO_3_ containing 0.5 % BSA. Dogs sera and secondary anti-dog IgG-HRP antibodies were used in 1:1,600 and 1:50,000 dilution, respectively. The statistically significant differences between exa groups are marked with an asterisk: *p < 0.05.

**Figure 11.**
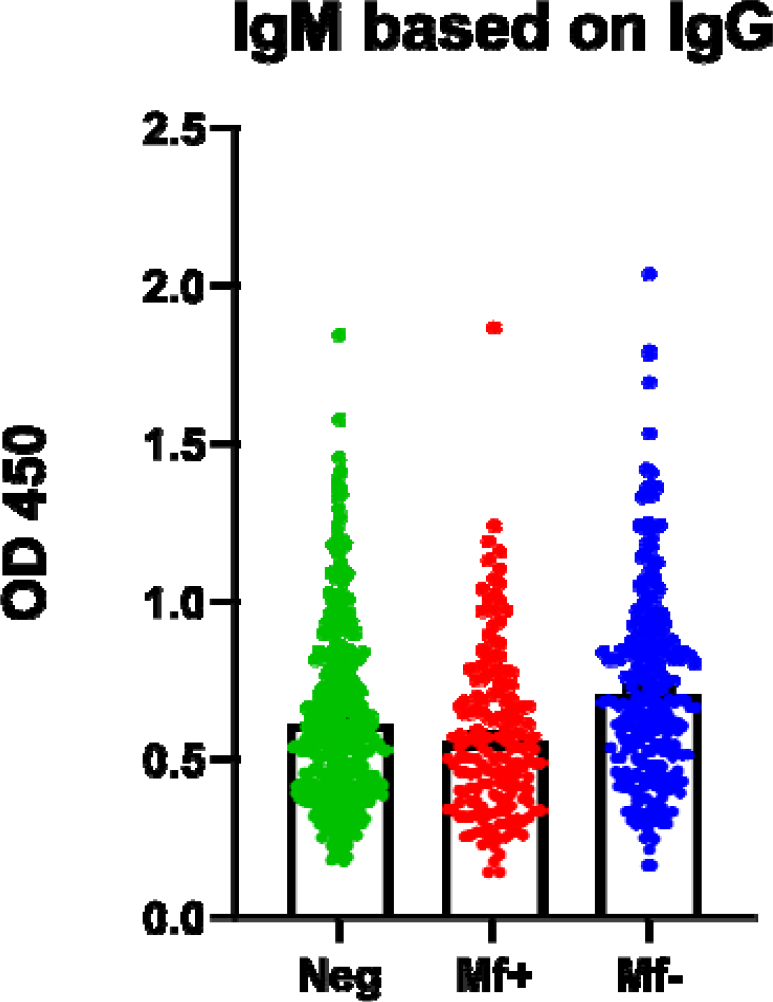
*D. repens* somatic antigen (DrSA) ELISA did not show statistically significant differences in IgM level between infected and non-infected groups. The IgM response to crude antigen from adult worms (DrSA) was evaluated in groups of dogs classified based on IgG level: microfilaremic(red N = 174) amicrofilaremic (blue N = 297) negative (green N =379) plates were coated with 2.5 µg/ml of DrSA in carbonate buffer. Dogs sera were used in 1:800 dilution in blocking buffer. Secondary anti-dog IgG-HRP antibodies were used in a dilution of 1:50,000 in blocking buffer.

**Figure 12.**
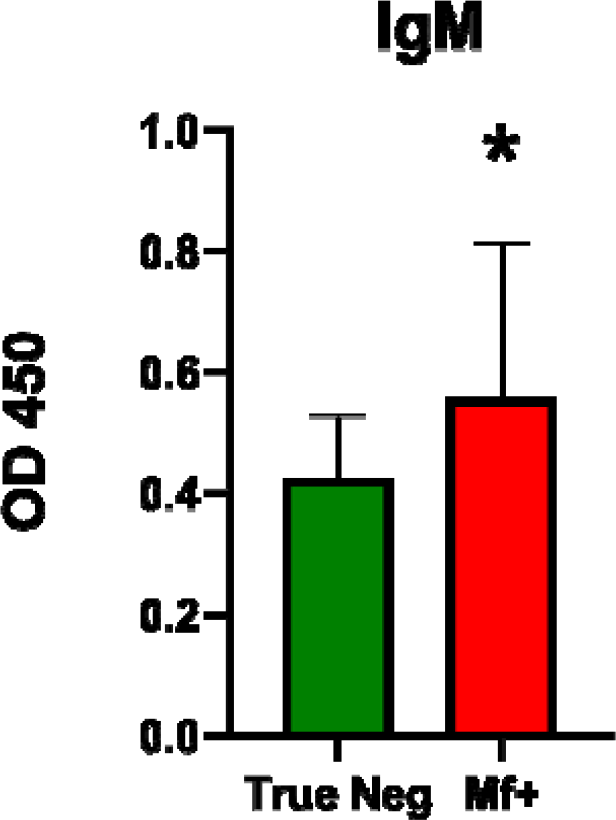
D. repens somatic antigen (DrSA) ELISA showed higher IgM level in microfilaremic dogs (red; N = 174) compared to “true negative” dogs (green; N = 208). Dogs included in the negative group present both low IgG and IgM levels. Plates were coated with 2.5 µg/ml of DrSA in carbonate buffer. Dogs sera were used in 1:800 dilution in blocking buffer. Secondary anti-dog IgG-HRP antibodies were used in a dilution of 1:50,000 in blocking buffer. The bars show mean ± SEM. The statistically significant differences between examined groups are marked w th an asterisk: *p < 0.05.

**Figure 13.**
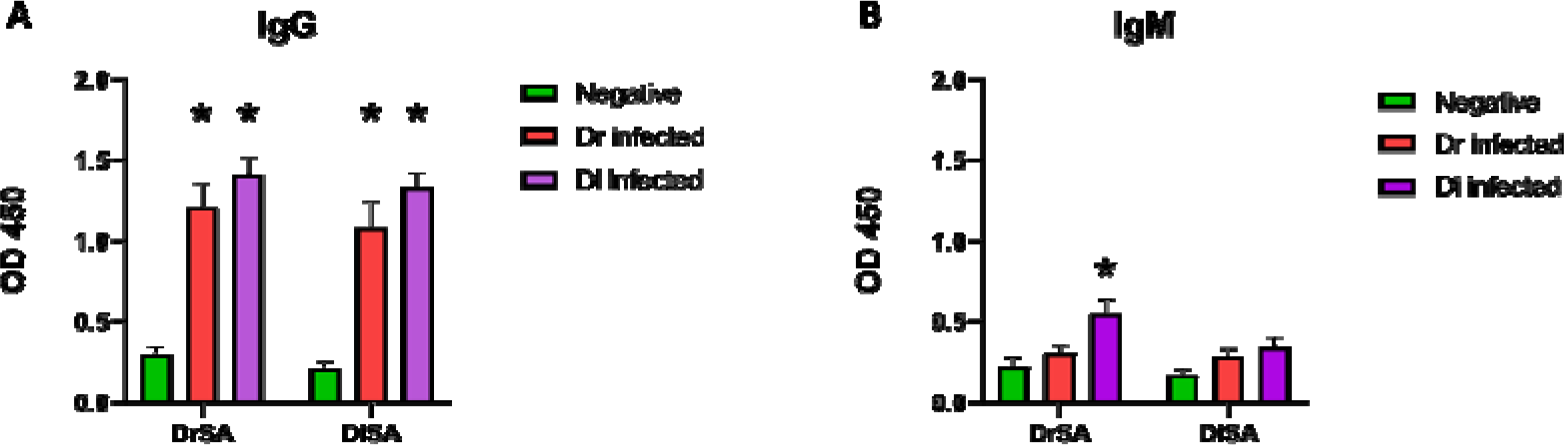
DrSA ELISA does not allow for differentiation between D. repens and D. immitis infections. Cross-reactivity in both IgG (A) and IgM (B) from dogs infected with either D. repens (red; N = 9) or D. immitis (purple; N = 9) was observed between crude antigen from adult worms of both species (DrSA vs. DiSA). Plates were coated with 2.5 µg/ml of DrSA/DiSA in carbonate buffer, then blocked with 0.1 M NaHCO3, 0.5 % BSA and washed three times. Dogs sera were used in 1:1,600 dilution in blocking buffer. After three washes, secondary anti-dog IgG-HRP antibodies were used in a dilution of 1:50,000 in blocking buffer. The bars show mean ± SEM. The statistically significant differences between examined groups are marked with an asterisk: *p < 0.05.

### 2.9. ELISA with Dr20/22 Reveals IgG Response in Both Infected and Negative Groups

We proposed that Dr20/22, similar to its homologues in other filariae species, could be a specific marker for infective larvae and a promising candidate for early diagnosis of subcutaneous dirofilariasis. After successfully purifying the protein in recombinant form and confirming the presence of specific IgG in sera from dogs infected with D. repens, we explored its potential as an alternative to using crude antigen from adult worms. We evaluated its diagnostic capabilities using sera from infected and healthy dogs and compared the results to the DrSA ELISA.

While the IgG levels were significantly higher in both infected groups, we noticed that many negative dogs also exhibited increased antibody levels (Figure 14). Given the limited detection of IgM antibodies against the protein (Figure 15), we decided to exclude the IgM class from further investigations.

**Figure 14.**
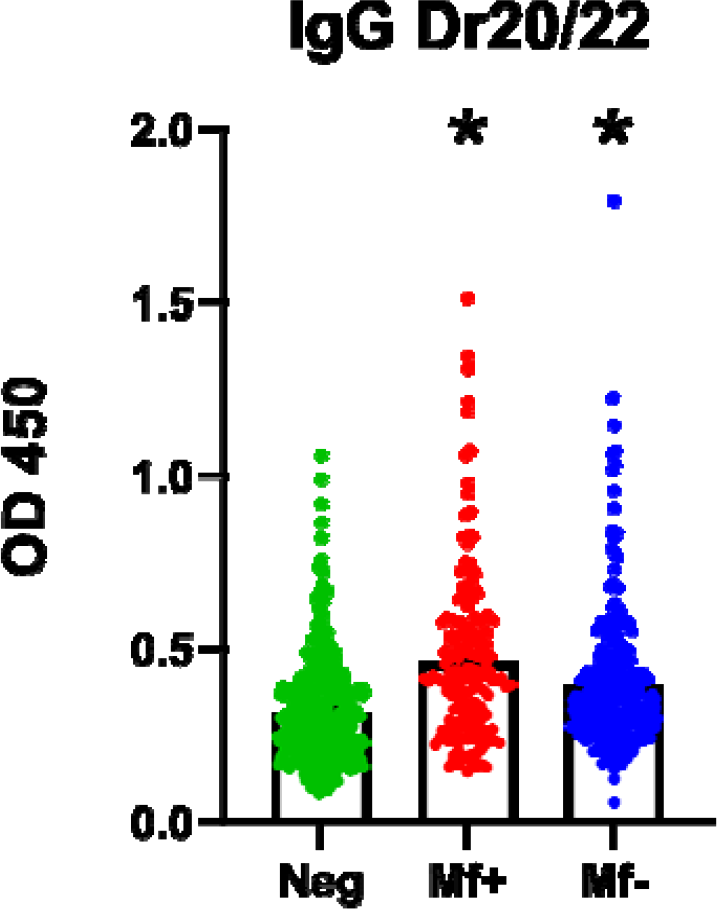
Dr20/22 ELISA demonstrated the presence of antigen-specific IgG in sera of both infected and healthy dogs. The IgG response to Dr20/22 was evaluated in groups of dogs classified according to the DrSA ELISA results: microfilaremic (red; N = 174), amicrofilaremic (blue; N = 297) and negative (green; N = 379). Plates were coated with 2.5 µg/ml of Dr20/22 in carbonate buffer. Plates were blocked with 0.1 M NaHCO3, 0.5 % BSA and washed three times. Dogs sera were used in 1:400 dilution in blocking buffer. After three washes, secondary anti-dog IgG-HRP antibodies were used in a dilution of 1:50,000 in blocking buffer. The statistically significant differences between examined groups are marked with an asterisk: *p < 0.05.

**Figure 15.**
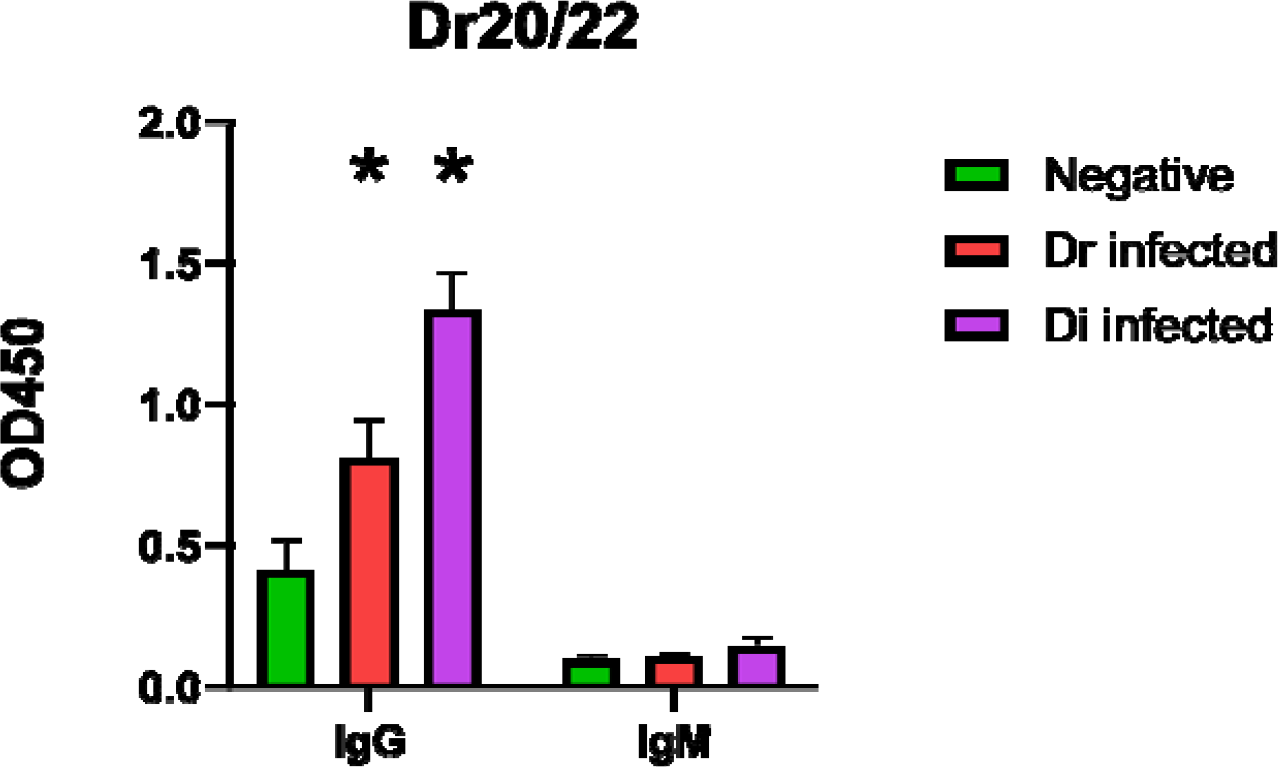
Comparison of the IgG and IgM responses to Dr20/22 in sera from healthy dogs (green; N = 3) and dogs infected with either *D. repens* (red; N = 9) or *D. immitis* (purple; N = 9). The level of IgM in both infected groups was barely detectable. Plates were coated with 2.5 µg/ml of Dr20/22 in carbonate buffer. Dogs sera were used in 1:400 dilution in blocking buffer. Secondary anti-dog IgG-HRP antibodies were used in a dilution of 1:50,000 in blocking buffer. The statistically significant differences between examined groups are marked with an asterisk: *p < 0.05.

Comparing the amino acid sequences of Dr20/22 and its homologue in D. immitis, we observed approximately 70 % identity, which is relatively low considering the high genetic similarity of both species. Furthermore, we identified a small deletion in positions 41-52 (Supplementary File S2), which might create a novel epitope in the protein. Interestingly, Western blot analysis detected specific IgG only in D. repens sera, not in D. immitis sera, leading us to consider the antigen’s potential in differentiating between the infections. However, ELISA results revealed that the antigen exhibited cross-reactivity with sera from both species.

## 3. Discussion

The study focused on ALT proteins (abundant larval transcript) found in filariae species, particularly their role in early diagnosis and immunogenicity. ALTs have been identified as high stage-specific proteins and are among the most abundant proteins in L3 larvae of several filariae species [10–16]. These antigens are stockpiled in the glandular esophagus in the form of inclusion bodies during larval growth in the mosquito and are later secreted via pseudocoelom and cuticle upon entry into the definitive host tissues, triggered by an increase in temperature [11,12].

In most species, there are at least two transcription variants (alt-1 and alt-2) sharing high amino acid identity (e.g., 79 % in *B. malayi*) but differing among homologues in the filariae family. For instance, in the case of *B. malayi, alt-2* and *alt-1* mRNAs are the first and third most abundant transcripts in the L3 stadium, accounting for 2.82 % and 1.39 % of all transcripts, respectively. However, in non-L3 stages, *alt* transcripts account for less than 0.01 % [11]. Additionally, *alt-1* expression in *B. malayi* is terminated abruptly just after injection into the mammalian host tissues, but *alt-2* expression is less rigorously controlled, and trace levels of the transcripts were observed in all life cycle stages. At the protein level, both variants of *B. malayi* ALTs are L3 specific, similar to its orthologs in D. immitis (Di20/22) [11,15,16].

In the present study, we successfully cloned cDNA and gDNA encoding the *D. repens* homologue of ALT, which we named Dr20/22. The predicted molecular weight was significantly lower than the actual size on the gel, similar to its orthologs in other filariae (15.1-15.8 kDa) [12,13]. The observed double band on the gel may be a result of posttranslational modifications, such as glycosylation and phosphorylation. *Brugia* ALT obtained in *Pichia pastoris* was also observed as a double band due to the glycosylation process [17]. Despite the presence of potential O-glycosylation sites, the glycan residues were not confirmed by PAS staining in the study. Computational analyses of the Dr20/22 predicted 18 phosphorylation sites. All the phosphorylation events could result in the addition of almost 1.5 kDa [26], which may explain the occurrence of the upper band of the Dr20/22.

Genomic cloning revealed that the dr20/22 gene is constructed with 4 exons and 3 introns (Supplementary File S1), which accords with the data from research on *B. malayi* alt genes. Gomez-Escobar et al. [27] identified two near identical copies of *alt-1 (alt1.1* and *alt1.2*) in the parasite genome, whereas alt-2 was shown to be encoded by a single polymorphic locus that may differ in the individual parasites. Interestingly, genes are not closely related to each other, and neither of the genes is formulated in an operon with any other gene. By analogy to other filariae, it can be expected that *D. repens* also has two distinct forms of ALT (Dr20/22). Based on the fact we observed expression of dr20/22 in both microfilaria and adult stages, we assume that we are dealing with ALT-2 like form.

In our study, we demonstrated that the expression level of dr20/22 mRNAs was approximately three times higher in the adult stage compared to microfilariae. However, surprisingly, we did not detect native protein in any of the tested *D. repens* lysates (female adult worm, adult worm E/S, microfilariae), indicating potential specificity of the antigen for L3 larvae. Furthermore, in immunohistochemistry, we observed a strong signal across almost the entire female cross-section, suggesting that the obtained anti-Dr20/22 serum may react non-specifically with other *D. repens* antigens, similar to what has been reported in the case of O. volvulus [12,28]. Although we were unable to directly compare the Dr20/22 expression level to L3 larvae due to the unavailability of L3 samples, our findings, in combination with available literature, lead us to propose that Dr20/22, like its orthologues, is specific to the infective larvae stage.

There are several potential explanations for why we were not able to detect the antigen in the studied stages. Firstly, it is possible that the expression level was too low to be detected using the described techniques. However, we reject this possibility, as in our previous study, we successfully identified Dre33 in microfilariae lysate, which presented a similar mRNA expression level to Dr20/22 [29]. The second, more plausible scenario involves molecular mechanisms responsible for the regulation of gene expression during larvae development. This phenomenon has been observed in research on *A. vitae* ES-62 antigen, where despite mRNA being expressed across the worm’s life cycle, the protein is only secreted by L4 and adult worms and has never been detected in the filarial stages. The authors considered the possibility of different mRNA stability, but they ruled it out as the 3’ UTRs were found to be identical in all stages [30]. Additionally, various regulatory elements, both constructed and unconstructed (linear), play a leading role in translation and are located in the 5’ UTR [31].

One such regulatory element is the spliced leader (SL) sequence, a short RNA containing a hypermodified 5’-cap structure that is transcribed from a different genomic location and transferred to the 5’ end of designated mRNAs in a process known as spliced leader trans-splicing (SLTS) [32]. SLTS was first described in 1982 in African trypanosomes in mRNAs encoding variant surface glycoproteins [33]. It has since been observed in several other phyla, including cestodes, trematodes, and nematodes [34–36]. In nematodes like *Ascaris lumbricoides* and *Caenorhabditis elegans*, the overall frequency of trans-spliced mRNAs ranges from 50 % to even 90 % depending on the source [37–40]. The exact mechanism of SLTS remains puzzling, but several distinct functions have been proposed, including providing a 5’-cap structure for protein-coding RNAs, resolving polycistronic into individual capped, monocistronic mRNAs, enhancing mRNA translational efficiency through the hypermodified cap structure and/or leader sequence, and trimming and sanitizing 5’ UTRs of the pre-mRNAs [32].

In most research on filarial ALTs, the antigens were cloned using an L3-derived RNA template and primers specific for the spliced leader (SL) sequence [10,12,13]. However, in the present study, we did not detect the SL in dr20/22 (*D. repens* homologue of alt) in any of the examined stages. Given the fact that we previously identified the same SL sequence (specific for all nematodes) in adult worm mRNA encoding the ES62 antigen, known to be an adult stage-specific antigen, we considered whether the SLTS in *D. repens* may be differently regulated throughout the parasite’s life cycle. Several studies suggest that leader sequence-containing transcripts may be stage-specific or developmentally regulated, which might account for differential protein levels and protein repertories in different stages [41–44]. As the parasite’s development is usually related to changes in the host/habitat, the involvement of environmental and physiological factors in controlling translation via SLTS cannot be excluded. For example, it has been shown that heat shock can inhibit the SLTS process in selected genes of trypanosomes [45]. In the study on the marine chordate *Oikopleura dioica*, it was observed that some of the SLs contain a TOP-like motif that may enable nutrient-dependent translation control of the trans-spliced mRNAs in response to changes in physiological and environmental conditions [46,47]. Similar trans-spliced TOP mRNAs have also been presented in *C. elegans*, suggesting that this phenomenon may occur in other species as well [46].

Despite the enigmatic nature of the precise mechanism of SLTS, it has been demonstrated that the process impacts translation efficiency [48–50]. For instance, a study on *A. lumbricoides* showed that the SL sequence and its hypermethylated cap enhance translational efficiency, especially at the initiation of protein synthesis [39]. Furthermore, Lall S. et al. [51] suggested that both structures may impact translation initiation of the trans-spliced transcripts due to trimming of primary transcripts that are non-optimal for translation. Interestingly, in the study on trypanosomes, the authors suggested that omitting SLTS/polyadenylation sites during polycistronic RNA processing may lead to the generation of RNAs in a “translational latency state”, which can be stored for further processing into a mature transcript when required by the cell [52].

To summarize, based on our findings and available data, we propose that *D. repens dr*20/22 gene expression may be developmentally controlled by the SLTS. We hypothesize that in vertebrate-specific stages (mf, adult), the gene is transcribed into non-functional mRNAs, which contain problematic elements in the 5’ UTR that prevent efficient translation. However, after the mosquito ingests microfilariae and the molting process starts, environmental or physiological changes promote SLTS events that sanitize alt’s 5’ UTR and add a modified cap structure, resulting in effective translation. After injection of L3 into the definitive host tissues, the temperature seems to signal abrupt secretion of the antigen and SLTS inhibition at once, leading to the presence of only “latent” transcripts in the adult stage. For a better understanding of the gene expression pattern and detailed mechanism, further study should be conducted using *D. repens* L3 larvae.

Although ALT proteins were identified over 20 years ago, their biological activity and processes in which they are involved are still unknown. Some studies have reported a weak similarity to phospholipase A2 enzymes (PLA2), which may be implicated in membrane homeostasis, remodeling, migration through host tissues, or signal transduction [13,15]. In our present study, we constructed the most probable 3D structure of the protein based on PLA2, but the confidence of the model reached less than 4 %. Finally, we excluded this hypothetical activity of Dr20/22 using a commercially available phospholipase A2 assay kit.

The second part of the study examined the diagnostic potential of the antigen. Despite its unknown biological function, the prophylactic potential of the antigen has been widely investigated, especially in human lymphatic filariasis caused by *B. malayi* and *W. bancrofti* [11,14,17,18,20]. Antibodies specific for the *D. immitis* homologue of ALT were also presented in sera from putatively immune dogs but not in infected, non-immune dogs [16]. In our study, we tested the diagnostic potential of Dr20/22 using ELISA and compared the results to ELISA with parasite somatic antigens [53–56]. The cut-off value calculated based on IgG response in true negative dogs allowed specific diagnosis and classification of dogs into three groups (microfilaremic, amicrofilaremic, negative). The analysis of IgM levels did not reveal any significant correlation between infected and non-infected groups, what stands in contrast to our recent study. However, it should be noted that all dogs included in the previous study were tested molecularly [25]. In the present study, we tested ~850 dogs in ELISA, so we were unable to confirm all of them using our standard protocol (Knott + ELISA + Real-Time PCR). No differences in IgM levels between infected and healthy dogs were also observed in a study conducted by Wysmołek et al. [57]. The authors considered the presence of natural IgM characterized by low affinity and non-specific binding to epitopes mimicking *D. repens* antigens as a potential explanation.

Additionally, we observed strong cross-reactions between sera from dogs infected with *D. repens* or *D. immitis* and DrSA / DiSA extracts. Consequently, while the ELISA allowed for specific diagnosis in the IgG class, it cannot be used to differentiate between these infections.

ELISA with Dr20/22 revealed the presence of specific IgG in both infected (mf+, mf-) and non-infected (negative) groups, similar to the findings in other filarial ALTs [11,12,14,18]. A number of dogs classified as negative based on DrSA presented increased IgG levels, indicating potential immunity to dirofilariasis. Similarly, a research on *Brugia* showed that the mean level of total IgG against ALT-2 in endemic normals (EN) and chronic pathology (CP) was even higher than in patients with active microfilaremia (MF) [17].

We did not test the antigen in the IgM class as the signal observed in the preliminary result (Figure 15) was barely detectable. Interestingly, despite cross-reactions observed between both *Dirofilaria* species in Dr20/22 ELISA, Western blotting specifically recognized the antigen only in dogs infected with *D. repens*. Similar results were observed in research on Di20/22 (*D. immitis* homologue of the ALT), which was not recognized by dogs infected with heartworm [16]. Moreover, in native protein analyses (Dot blot, ELISA), we noticed the protein was recognized not only by dogs infected with both *D. repens* and *D. immitis*, but even by non-infected (potentially immune) dogs. This may indicate that the antigen conformation (native vs. denatured) impacts its antigenicity. The 3D structure of Dr20/22 shares common epitopes for both parasites, but when the protein is denatured and unfolds, its epitopes may take a linear form and become specific only for *D. repens*. Although Dr20/22 may not be a perfect diagnostic candidate, it may still be useful in differentiating infections using Western blotting.

It has been shown that the depletion of antibodies specific for ALT from the sera samples of EN strongly declines the ability of the sera to participate in the elimination of *B. malayi* L3 in an ADCC assay [58]. This suggests that the acquisition of resistance against filarial diseases may be mediated by the presence of ALT-specific antibodies. Furthermore, as ALTs are highly expressed in invasive larvae and have no known homologue in mammals, they are considered one of the most attractive vaccine targets in filarial infections. Depending on the study and adopted vaccine strategy (single molecule vs. cocktail of molecules), the protection efficacy ranges from 30 % to 96 % and 70 % to 80 % using only *alt* DNA and *alt* DNA mixed with other constructs, respectively [21–24]. In the case of protein-based vaccines, the mean reduction of the alive larval burden reached 69 % to 76 % when using *B. malayi* ALTs [11,21–23], and 49.82 % to 62.26 % based on *W. bancrofti* recombinant ALT or ALT mutants (e.g. protein without acidic domain or signal peptide) [14]. These antigens provided better protection than any other previously tested filarial protein and reached efficacy that rivals vaccination with radiation-attenuated larvae [59].

In conclusion, our study provides valuable insights into the expression pattern and potential specificity of Dr20/22 in *D. repens*. It also highlights the role of SLTS in gene expression regulation during the parasite’s life cycle. However, further research with L3 larvae is needed for a better understanding of the gene expression pattern and its mechanisms. Additionally, the biological activity and processes involving ALT proteins remain unknown, and more studies are required to explore their potential role in vaccination and immunity against filarial infections.

## 4. Materials and Methods

### 4.1. Dr20/22 Cloning and Recombinant Protein Production

Total RNA was extracted from adult *D. repens* worms and microfilariae using the Total RNA Mini Kit (A&A Biotechnology), following the manufacturer’s protocol. The adult worms were manually homogenized in phenozol solution, while the microfilariae were isolated from blood using 5 µm filters (Whatman), suspended in phenozol solution, and homogenized in TissueLyser LT (Qiagen, Hilden, NRW, Germany). Each RNA sample (1 µg) was subjected to DNase treatment and cDNA synthesis using the RevertAid First Strand cDNA Synthesis Kit (Thermo Fisher Scientific) as per the manufacturer’s instructions, with the PTX primer (5’ GAA CTA GTC TCG AGT TTTTTTTTTTTTTTTTTT 3’) replacing oligo (dT)18. The adult worm cDNA mixture was used as a template in PCR with PTX and gene-specific ForDr20/22 (5’ ATG AAC AAA CTT YTM ATA RTY YTY GGC 3’) primers to amplify the sequence encoding the *D. repens* homologue of the ALT (Dr20/22). The gene-specific primer was designed based on homologous genes from closely related filariae species (GenBank: U29459.1, U47545.2, AF285860.1). The PCR reaction was performed under the following conditions: 3 min at 95 °C, 35 cycles of 95 °C for 30 s, 53 °C for 45 s, and 72 °C for 30 s, with a final extension step of 10 min at 72 °C. The full-length cDNA was ligated into the pGEM T-Easy vector (Promega) and sequenced using Sanger’s method to confirm the specificity of the product. Additionally, PCR was performed with ForDrSL (5’ GTT TTA ATT ACC CAA GTT TGA GG 3’) and gene-specific RevDr20/22cds (5’ ATC ATA TGA GCA CTG CCA GTC 3’) primers to test if the mature adult and microfilariae mRNAs are enriched with the spliced leader (SL) sequence specific for *D. repens*, as detected in a previous study (GenBank: MT071086.1). The reaction conditions were the same as described above.

Subsequently, cDNA encoding the mature protein (without the signal peptide) was amplified using gene-specific primers incorporating restriction enzyme sites and subcloned into the pPICZα A vector (Invitrogen) using EcoRI and XbaI enzymes and T4 ligase (Thermo Fisher Scientific). The resulting construct (10 µg) was linearized with SacI enzyme and used for electrotransformation of the *P. pastoris* X33 strain. Transfected cells were spread on YPDS plates with increasing concentrations of zeocin (100, 500, 1000 µg/ml) to select multi-copy recombinants. A single colony positive in PCR with 5AOX1 (5’ GAC TGG TTC CAA TTG ACA AGC 3’) and 3AOX1 (5’ GCA AAT GGC ATT CTG ACA TCC 3’) primers was used for expression in BMMY medium containing 0.5 % methanol at 29.8 °C for 72 h. All yeast transformation, culture, transformant analysis, and protein expression procedures were performed according to the EasySelect™ Pichia Expression Kit (Invitrogen) protocols.

The protein was purified under native conditions using the Protino Ni-NTA Agarose for His-tag protein purification Kit (Macherey-Nagel) with the gravity flow method, following the manufacturer’s protocol. Eluted fractions were concentrated and dialyzed against PBS using Amicon Ultra-15 Centrifugal Filter Units with a 3 kDa MWCO (Merck Millipore). Endotoxins were removed using High Capacity Endotoxin Removal Spin Columns (Pierce), and the purified Dr20/22 protein was filtered through 0.22 µM syringe filters (Millex). The protein concentration was determined using the BCA Protein Kit Assay (Pierce). The presence of purified Dr20/22 was confirmed by SDS-PAGE and Western blotting using the Anti-polyHistidine Peroxidase antibody (Sigma). The occurrence of potential glycosylation sites was assessed using the Glycoprotein Staining Kit (Pierce).

### 4.2. Cloning of Dr20/22 gDNA

Adult *D. repens* worms were manually homogenized in tris buffer, and genomic DNA (gDNA) was isolated using the Genomic Mini Kit (A&A Biotechnology) following the manufacturer’s protocol. The isolated gDNA was utilized as a template in PCR with primers designed to target the start (ForDr20/22cds 5’ ATG AAC AAA CTT TTC ATA GTT CTT GGC 3’) and stop (RevDr20/22cds 5’ ATC ATA TGA GCA CTG CCA GTC 3’) codons of the coding sequence. This PCR amplification aimed to capture the full-length gene, including both exons and introns. The reaction conditions involved an initial denaturation at 95 °C for 3 min, followed by 35 cycles of denaturation at 95 °C for 30 s, annealing at 52 °C for 45 s, and extension at 72 °C for 2 min, with a final extension step of 10 min at 72 °C. The amplified product was then cloned into the pGEM T-Easy Vector (Promega) and subjected to Sanger sequencing. The obtained sequence data were subsequently used for the prediction of the gene structure, including the determination of the number of exons and introns.

### 4.3. Analysis of Gene Expression Level

Gene expression levels were assessed using Real-Time PCR. Total RNA isolation, DNase treatment, and cDNA synthesis were performed following the previously described methods. PCR reactions were conducted in triplicates, employing a two-step fast cycle protocol with PowerUp™ SYBR™ Green Master Mix from Applied Biosystems™. The melt curve step was carried out in a QuantStudio 6 Real-Time PCR system (Applied Biosystems). The PCR process involved UDG (Uracil-DNA Glycosylase) activation for 2 min at 50 °C, followed by initial denaturation for 2 min at 95 °C. Subsequently, 40 amplification cycles were executed, comprising 3 s at 95 °C and 30 s at 60 °C. The reaction mixture included 10 ng of cDNA from each life cycle stage (adult *D. repens* worm and microfilariae), 5 µl of PowerUp™ SYBR™ Green Master Mix (2X), forward (ForDr20/22RT 5’ CAG CGA CGA AAG TTA TGC AGA AGA C 3’) and reverse (RevDr20/22RT 5’ AAT ATG CAC CAC GAT TGC GGT TCA C 3’) primers at a final concentration of 0.6 µM, and sterile water to reach a final volume of 10 µl. The absolute copy number of the dr20/22 gene was determined based on a standard curve ranging from one billion to ten copies per reaction, utilizing the data collected during the anneal/extension step.

Statistically significant differences between different life cycle stages were analyzed using the Student’s t-test.

### 4.4. Bioinformatics Analysis and Structural Prediction of D. repens Dr20/22 Protein

Comparison of the nucleotide sequences of alt from closely related filariae in the GenBank database was performed using the Basic Local Alignment Search Tool (Blast) available at http://www.ncbi.nlm.nih.gov/blast/Blast.cgi. The amino acid sequence was obtained using the Translate tool available at http://web.expasy.org/translate/. Nucleotide and amino acid alignments, incorporating the novel *D. repens* Dr20/22 sequence and homologous sequences from related filariae, were constructed using Multalin accessible at http://multalin.toulouse.inra.fr/multalin/. The theoretical molecular weight and isoelectric point were estimated using the Compute pI/Mw tool (https://web.expasy.org/compute_pi/). The signal peptide was determined using the SignalP 5.0 server available at https://services.healthtech.dtu.dk/services/SignalP-5.0/. For the determination of N-glycosylation, O-glycosylation, and phosphorylation sites, NetOGlyc, NetNGlyc, and NetPhos, respectively, were employed and accessed at www.cbs.dtu.dk/services. The identification of characteristic domains was carried out using the InterProScan Sequence Search available at https://www.ebi.ac.uk/interpro/. Lastly, the probable three-dimensional structure of *D. repens* Dr20/22 was obtained and visualized using Phyre2, accessible at http://www.sbg.bio.ic.ac.uk/phyre2/.

### 4.5. Dirofilaria E/S Collection and Tissue Lysates Preparation

Briefly, to collect D. repens excretory/secretory (E/S) products, several live worms obtained during surgical procedures were washed in PBS and then incubated in RPMI 1640 supplemented with penicillin (100 U/ml) and streptomycin (100 µg/ml) at 37 °C for 16 h. Fresh medium was replaced every 2 h, and the collected medium was stored at -70 °C until further analyses. Subsequently, the collected medium was centrifuged (8,000 × g, 10 min, 4 °C), concentrated, and dialyzed against cold sterile PBS using an Amicon Ultra-15 Centrifugal Filter Unit with a 3 kDa MWCO (Merck Millipore). The resulting product was then passed through a 0.22 µM filter.

For adult worm (D. repens and D. immitis) and microfilariae lysates, the procedures described in our previous studies were followed [25,29]. Protein concentration in the lysates was determined by BCA assay (Pierce).

### 4.6. Blood Sample Collection

All blood samples from dogs were collected and stored following the procedures detailed in our previous study [25].

### 4.7. Preparation of Mouse Anti-Dr20/22 Serum and Western Blotting Analysis of D. repens Proteins

Antiserum against recombinant Dr20/22 was generated through subcutaneous immunization of a BALB/c mouse with 100 µg of the recombinant protein, followed by three booster doses of 75, 50, and 25 µg on days 14, 28, and 42, respectively. Each dose was mixed with Imject™ Alum Adjuvant (Thermo Fisher Scientific) in a 1:3 ratio. On day 49, the mouse was euthanized, and blood was collected in tubes, incubated for 2 h at 37 °C, followed by overnight incubation at 4 °C. The next day, the blood was centrifuged for 10 min at 500 × g, and the serum was stored at -70 °C until further analyses. All experiments were conducted following relevant guidelines and regulations, and ethical approval was obtained from the 2nd Local Ethics Committee for Animal Experimentation, Warsaw University of Life Sciences-SGGW, Poland (approval number: WAW2/82/2019). The manuscript reporting adheres to the recommendations in the ARRIVE guidelines [60].

To determine the presence of antibodies recognizing recombinant Dr20/22 epitopes in the mouse serum and to detect native protein in adult worm lysate, excretory/secretory (ES) products, and microfilariae lysate, Western blot analysis was performed. 0.25 µg of the recombinant protein and 10 µg of each lysate were separated on a 15 % polyacrylamide gel during SDS-PAGE and subsequently transferred to a nitrocellulose membrane. The membrane was blocked for 2 h at room temperature using SuperBlock T20 (Thermo Fisher Scientific) and then incubated overnight at 4 °C with mouse anti-Dr20/22 serum or serum from non-immunized mice, both diluted 1:1000 in the blocking buffer. The following day, the membrane was thoroughly washed with PBS containing 0.05 % Tween-20 (0.05 % PBS-T) and then incubated for 1 h at room temperature with Anti-Mouse IgG HRP-conjugated antibodies (R&D Systems), diluted 1:1000 in the blocking buffer. After washing with 0.05 % PBS-T (3 × 10 min), the membrane was visualized by chemiluminescence using SuperSignal™ West Pico PLUS Chemiluminescent Substrate (Thermo Fisher Scientific) in a Chemidoc MP Imaging System (Bio-rad).

### 4.8. Immunohistochemical Detection of Dr20/22 Protein in Adult Female D. repens Worm

Detection of the Dr20/22 protein in adult female *D. repens* worms was carried out using immunohistochemistry staining. The worms were fixed in 4 % buffered formalin, embedded in paraffin, and cross-sectioned to a thickness of 5 µm using a Leica RM2025 microtome (Leica Microsystems, Germany). The obtained histological sections were initially subjected to topographic staining using the hematoxylin-eosin (H/E) standard protocol to assess the morphology of the adult nematode.

For the immunohistochemistry staining, the histological slides were deparaffinized in xylene, rehydrated through a gradient of ethanol (from absolute alcohol to 70 %), and then subjected to antigen retrieval by incubation for 5 min at 82 °C in 10 mM citric acid (pH 6). Endogenous peroxidase was blocked using 5 % hydrogen peroxide, and to reduce non-specific antibody binding, the slides were blocked with Protein Blocker (Novocastra Peroxidase Detection System, Leica) and 5 % skim milk, each step lasting 30 min at 37 °C. Subsequently, the slides were incubated with polyclonal anti-Dr20/22 mouse serum, diluted 1:100 in PBS, overnight at 4 °C. The following day, incubation with secondary antibodies and visualization were performed according to the manufacturer’s protocol (Novocastra Peroxidase Detection System, Leica). After each step, the slides were rinsed in tris buffer (pH 8.0, Sigma). Finally, the slides were counterstained with Harris hematoxylin solution, differentiated with 1 % acid alcohol, dehydrated, rinsed in xylene, and mounted in DPX Mountant for histology (Sigma). For the negative control test, sera from non-immunized mice were used.

### 4.9. Immunological Analysis of Dr20/22 Antibodies in Dogs Infected with D. repens or D. immitis

Western blot analysis was performed to determine the presence of antibodies specific for Dr20/22 in sera from dogs infected with *D. repens* or *D. immitis*. The membrane was incubated with sera from infected and healthy dogs, diluted 1:400 in the blocking buffer, overnight at 4 °C. Subsequently, secondary anti-dog IgG-HRP antibodies (abcam) were incubated for 1 h at room temperature in a 1:25,000 dilution. The remaining procedure steps, including protein separation on the gel, washing, and visualization, were conducted as described in Section 4.7 [*Preparation of Mouse Anti-Dr20/22 Serum and Western Blotting Analysis of D. repens Proteins*].

In addition, Dot blot analysis was performed to evaluate the impact of the protein conformation on its antigenicity. 0.5 µg of Dr20/22 was deposited in several replicates on the nitrocellulose membrane and then blocked, washed, and incubated with sera from infected and healthy dogs as described above.

### 4.10. Comparative Diagnostic Evaluation of Dr20/22 ELISA and Crude Adult Worm Antigens in Canine Dirofilariasis

The Nunc MaxiSorp™ C-shaped Plates (Thermo Fisher) were coated with 2.5 µg/ml of DrSA/DiSA in 0.1 M sodium carbonate buffer (pH 9.5) and incubated overnight at 4 °C. Subsequently, the plates were blocked with 0.1 M NaHCO3 (pH 8.6) and 0.5 % BSA for 1.5 h at room temperature. Plasma samples from all subjects were analyzed in a dilution of 1:1,600 in the blocking buffer for 1.5 h at room temperature. The anti-dog peroxidase-conjugated IgG and IgM (abcam) were diluted to 1:50,000 in 0.1 M NaHCO3 (pH 8.6) and 0.5 % BSA and then incubated for 1 h at room temperature. After each step, the plates were washed three times with PBS containing 0.05 % Tween-20. TMB substrate solution was added to the wells, developed at room temperature, and the reaction was stopped after 30 min with 2 M H2SO4. Optical densities were measured at 450 nm using a microplate reader (Synergy HT, BioTek).

The cut-off value was calculated as the mean plus 3 standard deviations (SD) of 35 sera from dogs negative in Knott and qPCR, based on the previous study protocol [25].

ELISA with Dr20/22 was carried out to compare the diagnostic potential of the antigen to the crude adult worm antigens. All procedures were performed as described above, with the exception of the dogs’ sera, which were used in a dilution of 1:400. ELISA with Dr20/22 was carried out to compare diagnostic potential of the antigen to the crude adult worm antigens. All procedures were performed as described above with the exception of the dogs sera which were used in 1:400 dilution.

### 4.11. Evaluating the Phospholipase A2 (PLA2) Activity of Dr20/22

The potential phospholipase A2 (PLA2) activity of Dr20/22 was evaluated using the EnzChek™ Phospholipase A2 Assay Kit (Invitrogen). Fresh or frozen (stored at -80 °C) protein was utilized in two-fold dilutions ranging from 100 µg per reaction to 0.2 µg per reaction. Additionally, the ability of Dr20/22 to inhibit PLA2 activity was investigated. Each well was supplemented with 1.25 U/ml of PLA2, which was pre-incubated for 20 min with serial dilutions of Dr20/22 (ranging from 100 µg to 0.2 µg per reaction) before the substrate-liposome mix was added. All reactions were conducted in a 96-Well Black Polystyrene Microplate (Costar) following the manufacturer’s protocol.

The fluorescence was measured with excitation at ~485/20 nm and fluorescence emission at ~528/20 nm using a microplate reader (Synergy HT, BioTek). The PLA2 activity was determined based on a standard curve ranging from 5 U/ml to 0.2 U/ml.

### 4.12. Statistical Analysis

All data are presented as mean ± standard error of the mean (SEM). Statistical analysis was performed using GraphPad Prism 8.0 (GraphPad Software, La Jolla, CA, USA) with unpaired t-test for gene expression analysis and U Mann-Whitney test or Kruskal-Wallis test for data produced in ELISA. Differences between groups were considered statistically significant at p < 0.05.

## Supplementary Materials

The following supporting information can be downloaded at: (…), Supplementary File S1: Genomic DNA sequence of dr20/22 with determined exons and introns.; Supplementary File S2: Comparison of amino acid sequences of D. repens ALT and two variants of *D. immitis* ALT.

## Author Contributions

Conceptualization, M.P., M.W. and A.Z.-D.; methodology, M.P., M.W., and A.Z.-D.; software, M.P. and K.B.; validation, M.P., K.B., M.K., .; formal analysis, M.P., K.B.; investigation, M.P., K.B., M.K.; resources, M.P., E.C. and R.M.; data curation, M.P.; writing—original draft preparation, M.P.; writing—review and editing, M.P., K.B., D.M., M.W. and A.Z.-D.; visualization, M.P. and A.Z.-D.; supervision, M.W. and A.Z.-D.; project administration, A.Z.-D. All authors have read and agreed to the published version of the manuscript.

## Funding

This research was funded by National Centre for Research and Development, grant number 0106/L-9/2017.

## Institutional Review Board Statement

Ethical review and approval were waived for this study since all analyses were conducted using the leftovers of blood samples obtained from dogs during routine checkups or diagnostic procedures conducted by veterinarians at the veterinary clinic. These actions were performed in accordance with appropriate guidelines and regulations. Written consent was not acquired from the owners, as no interventions other than standard care were conducted. The results of the Dirofilaria testing were communicated to the dog owners.

## Informed Consent Statement

Not applicable.

## Data Availability Statement

The data presented in this study are available on request from the corresponding author.

## Acknowledgments

Not applicable.

## Conflicts of Interest

The authors declare no conflict of interest.

## Notes

### Competing Interest Statement

The authors have declared no competing interest.

