## Supplementary File S1 for "Characterization of Dr20/22: Molecular Insights and Antibody Response in Naturally Infected Dogs with *Dirofilaria repens*"

Genomic DNA sequence of *dr20/22* with determined exons and introns.

ATGAACAAACTTTTCATAGTTCTTGGCTTAGTGATTCTTTCTGTTGCATTACCTTCTGCATCAGAATCAGAAGAAGAGGTAATTTCTTCAGTTCAGTTTCTCAACTTTTAATAATATCAAATTAAAAGTAATCAGCAACAGGAATAATGACCAAATAAGGAAGATAAGGAATATATATTGAAGAGAATACATGCTAATCAAGTGCACTACTTCTTGGATTGGAGAAAAAGAAAAAGATTATGTTCAAGTCATTTTTTCGATCAAAATTTTCATAAAAGTATCTTTATGATTTCAGAGTGTATCTTTTGAAGACAGCGACGAAAGTTATACAGAAGATGATGAAGGTCATAAAGAAGAACACAATGATCATGCAACTGAAGACGATGAATATGTAACTAAAGGAAAATTTGTTGAAAGTGATGGTAAGTAAGTCGTTTCAATCTTTCAAGTATTTAAATTCCTGAAAACTTCATTAAAACTTTAGTTCGCAACCATATGTGTTCTTCTATCCATCCTGCAGCGCTGTTCCGCTACTCGAAATATAATTTCGCTATAATATGTTTTAAATTTCTTCATTTAAATAATGCGATTCAATAACTTTCCCAATACATTACAGATGATTTCCAATATGGACTGATTCGCTTACATTTCATTTTAATTTCTTGCAGAATGAAACACTGCAAAACTCATGAAGCTTGCTATGACCAACGTGAACCGCAATCGTGGTGCATATTGAAACCGCATCAATCATGGACGTAAGCACCTTATGATTATTCTAACGAAAAAAATAGTTTAGATAAGAATTTTCGATGCCTATTAATCAGAATTTCCGATTCCTATAAGTCAGAATTTCCTTCTAAATCAATATCCATAAATGTCTATAAAATCCATAAATTTCGCAATAAAATTGAGAACTTTTTGTCTTAGAAAAAGAGGTTGTTTCTGCGAATCAAAAAAGCATGCATGCGTTATCGAGCGGAAGAGCGGCGACAACTTTGAATATTCATATTGCTCACCACGAAAAGACTGGCAGTGCTCATATGATTAA

Exon1: 1-78 bp **(78 bp)**

Exon2: 296 - 422 bp **(127 bp)**

Exon3: 665 -755 bp **(91 bp)**

Exon4: 928 - 1,048 bp **(121 bp)**

STOP CODON
