## Supplementary File S2 for "Characterization of Dr20/22: Molecular Insights and Antibody Response in Naturally Infected Dogs with *Dirofilaria repens*"

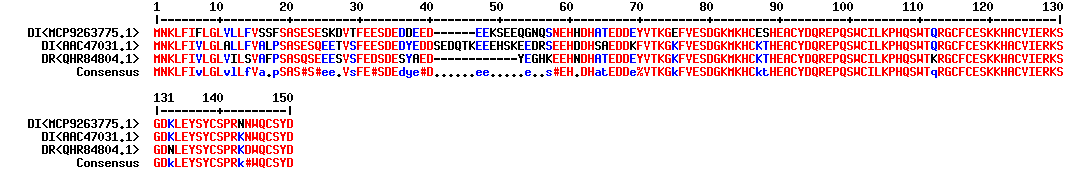

Supplementary File S2 Comparison of amino acid seqences of *D. repens* ALT (QHR84804.1) and two variants of *D. immitis* ALT (MCP9263775.1; AAC47031.1).
